# Validating molecular markers for barley leaf rust resistance genes *Rph20* and *Rph24*

**DOI:** 10.1101/2020.07.12.199075

**Authors:** PM Dracatos, RF Park, D Singh

## Abstract

Improving resistance to barley leaf rust (caused by *Puccinia hordei*) is an important breeding objective in most barley growing regions worldwide. The development and subsequent utilisation of high-throughput PCR-based co-dominant molecular markers remains an effective approach to select genotypes with multiple effective resistance genes, permitting efficient gene deployment and stewardship. The genes *Rph20* and *Rph24* confer widely effective adult plant resistance (APR) to leaf rust, are common in European and Australian barley germplasm (often in combination), and act interactively to confer high levels of resistance (Dracatos et al. 2015; Zeims et al. 2017; Singh et al. 2018). Here we report on the development and validation of co-dominant insertion-deletion (indel) based PCR markers that are highly predictive for the *Rph20* and *Rph24* resistances.

---

Leaf rust, caused by *Puccinia hordei*, is a serious foliar disease affecting barley world-wide. It has been reported to reduce yield by up to 62% in highly susceptible barley cultivars and the spread of the disease is usually exacerbated where monocultures of susceptible varieties dominate the landscape (Cotterill et al. 1992). Genetic resistance is the most desirable strategy to control leaf rust due to low environmental impact and cost effectiveness (Park et al. 2015). Although genetic resistance has been used to great effect, rapid evolution in pathogen populations often renders ineffective widely deployed resistance genes, especially in the case of single all-stage (major) resistance genes (Park 2003). In contrast to major resistance genes, partial resistance or APR is often more durable and is mainly conferred by multiple favourable alleles spread across the genome that often act additively (Niks et al. 2015).

To date, three genes conferring APR to *P. hordei* in barley have been catalogued based on their consistent detection and map location over multiple seasons in segregating mapping populations. *Rph20, Rph24* and *Rph23* confer high, moderate and low levels of APR respectively. Minor effect QTLs for partial resistance to BLR (*Rphq2, Rphq3* and *Rphq4*) were previously identified by Qi et al. (1998) in the Vada x L94 mapping population. *Rphq2* has been recently cloned and is involved in increased latency period, but is largely ineffective at adult growth stages (Wang et al. 2020). *Rphq4* and *Rph20* are regarded as the same major effect QTL and *Rphq3* is assumed to be the same as *Rph24* however further genetic evidence is required for confirmation. Hickey et al. (2011) mapped two QTLs on 5H and 6H in a doubled haploid (DH) population based on the cross ND24260 (*rph20*/*Rph24*) x Flagship (*Rph20*/*rph24*), and formally designated the *Rphq4* QTL on 5H from Flagship as *Rph20*. Recent studies also identified the *Rph20* resistance in mapping populations derived from European parental lines including Pompadour, Dash and Baronesse and the Australian barley breeding line WI 3407 using DArT-Seq complexity reduction markers (Singh et al. 2017; Dracatos et al. 2019). Zeims et al. (2017) mendelised the 6H QTL in an F2 population based on the cross ND24260 x Gus using the DArT-Seq platform and the QTL was subsequently designated as *Rph24*. The same QTL was mapped by Elmansour (2016) in an F3 population based on the cross CPI 36396A x Gus. Four highly significant DArT-Seq markers were highly predictive (Ziems et al. 2017), and determined that *Rph24* was present at high frequency (approx. 50%) in a diverse set of global germplasm (Singh et al. 2018).

Prospects for marker assisted selection (MAS) for the *Rph20* and *Rph24* resistance in global barley breeding programs is currently limited by a lack of co-dominant PCR-based markers. The current *Rph20* marker developed from DArT marker pBp-0837 (Hickey et al. 2011), although predictive, is a dominant marker type, and no markers are available for *Rph24*. We previously reported on the genetic characterisation of the minor APR gene *Rph23* including the validation of a closely linked (0.01cM) codominant SSR marker that is highly suitable for MAS (Singh et al. 2015). While co-dominant marker systems such as SSRs and single nucleotide polymorphisms (SNPs) have numerous benefits (multiallelic, elimination of false negatives etc), both also have limitiations; SSRs are found less frequently than SNPs, and SNPs are more expensive to assay. Although not as frequently identified as SNPs, indel events represent a low cost alternative to develop co-dominant PCR-based markers for efficient MAS.

In this study, we report the development and validation of co-dominant PCR-based indel markers to improve the efficiency of MAS for the *Rph20* and *Rph24* resistances. To identify indels and design PCR amplicons for size discrimination on agarose gels, we used the most strongly associated DArT-Seq marker from mapping studies for *Rph20* in a Recombinant Inbred Line (RIL) population based on the cross Pompadour x Biosaline-19 (Dracatos et al. 2019) and *Rph24* in an F3 population based on the cross ND24260 x Gus (Zeims et al. 2017). For example, for *Rph20* we used the 65bp DArT-Seq clone sequence from 3270804 as a BLASTn query to identify a homologous sequence from the Morex reference genome (http://webblast.ipk-gatersleben.de/barley_ibsc/). The highest Morex contig hit was then used as a BLASTn query against the Barke and Bowman assemblies, and where possible all three homologous contigs were aligned using Geneious V9.0.5 and the alignment was screened for the presence of indel events (Figure 1). For *Rph24*, the same approach was used to search for indels using the most closely associated DArT-Seq marker 3999875 from Zeims et al. (2017). Primer pairs for *Rph20* (*sun690* 5’ F AACAAAAAGCGGCCGAAAAA 3’ and *sun691* 5’ R ACGGGCACATTGTGTCTATTT 3’) and *Rph24* (*sun43* 5’ F CTAGACACCACCACCACACC 3’ and *sun44* 5’ R ATACCAGAGTTTGCGTCCGG 3’) were designed to flanking sequences to ensure the indel was at least 10% of the PCR product size to permit visualisation using gel electrophoresis. Optimal primer binding conditions were initially determined using a gradient PCR with annealing temperatures ranging from 55-70°C. The optimal annealing temperatures for indel markers closely linked to the *Rph20* (*sun690-91*) and *Rph24* (*sun43-44*) resistances were 55°C and 65°C, respectively. PCRs were performed using MyFi™ DNA Polymerase (Bioline), using 100ng of genomic DNA as described in the manufacturers instructions however an extension time of 20s was used for *sun690-91* relative to 30s for *sun43-44* due to differences in expected PCR product size.

**Figure 1.**
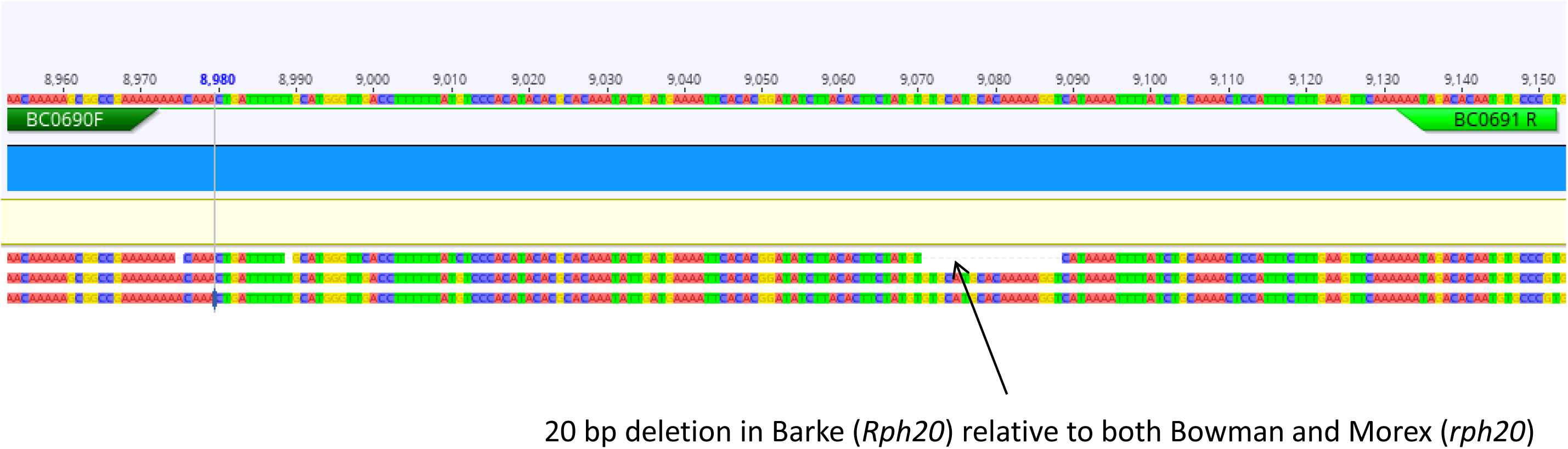
Schematic nucleotide alignment of contigs from barley cultivars Barke (*Rph20*), Morex (*rph20*), Bowman (*rph20*) with highest similarity match using the BLASTn algorithm in the https://webblast.ipk-gatersleben.de/barley_ibsc/ database to DArT-Seq clone 3270804 using Geneious v9.0.5. Primer pairs were designed to conserved sequence flanking the 5’ and 3’ of the 20 bp deletion in Barke relative to both Morex and Bowman.

For *Rph20*, a 20 bp deletion was found in Barke (*Rph20*) relative to the leaf rust susceptible accessions Bowman and Morex which are both susceptible to leaf rust. Interestingly, BLASTn analysis using both the 2017 (version 1) and 2019 (version 2) Morex reference genome interations indicated that DArT-Seq marker 3270804 had the strongest match to chromosome 4H, however linkage and QTL mapping in the Pompadour x B-19 RIL population (Dracatos et al. 2019) and three other populations mapped 3270804 on chromosome 5H co-locating with the *Rph20* resistance. This suggests either a problem in assembly or more likely pan-genomic variation between Morex and *Rph20*-carrying accessions such as Barke, Pompadour, Flagship and Vada. In contrast, the *Rph24* DArT-Seq marker 3999875 had a closest BLASTn match in the 2019 Morex assembly to 6H however, in the earlier 2017 assembly 3999875 had no strong similarity to any Morex chromosome.

Both primer pairs flanking the indels identified in contig assemblies at DArT-Seq loci 3270804 and 3999875 for *Rph20* and *Rph24* were validated on a collection of Australian barley cultivars with consensus disease ratings based on field data across multiple years and environments. We genotyped 115 Australian barley cultivars with indel markers for *Rph20* and *Rph24* that have been 1/phenotyped over multiple seasons and years in the field and 2/multipathotype tested in the greenhouse with *P. hordei* pathotypes with contrasting pathogenicity (Table 1). There was high correlation with the presence of APR and the marker alleles for *Rph20, Rph24*, or both in the case of highly resistant lines. Amongst the Australian barley varieties assessed, 56 (46%) carried the *Rph24* marker allele, almost twice the frequency of *Rph20* (25%). Interestingly, 75% of the lines that carried the *Rph20* marker allele also carried *Rph24* and this was characterised by a low consensus disease rating. Twenty Australian cultivars carried the marker alleles for both *Rph20* and *Rph24*, including: Chieftain, Capstan, Cosmic, Henley, Oxford, Quasar, Quickstar, Shepherd, Starmalt, Westminister and Dove. The combination of all three APR genes was however, not identified. Over half of these accessions postulated to carry both APR genes were from European decent, reflecting the strong influence of European germplasm exchange in Australian barley breeding programs. Representative genotypic results for the resistant and susceptibility alleles for indel markers linked to APR genes *Rph20* and *Rph24* are shown in Figure 2.

**Table 1.**
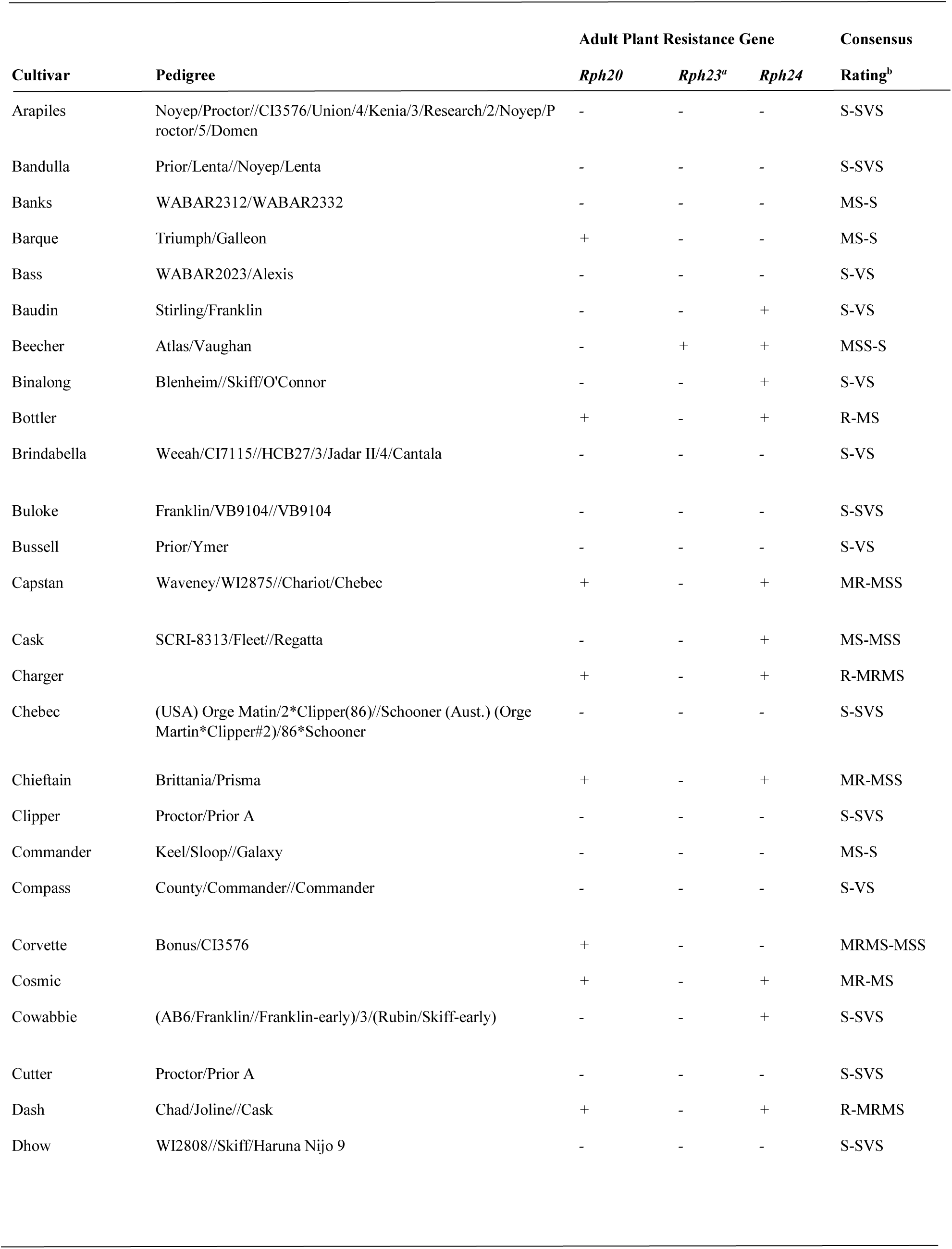
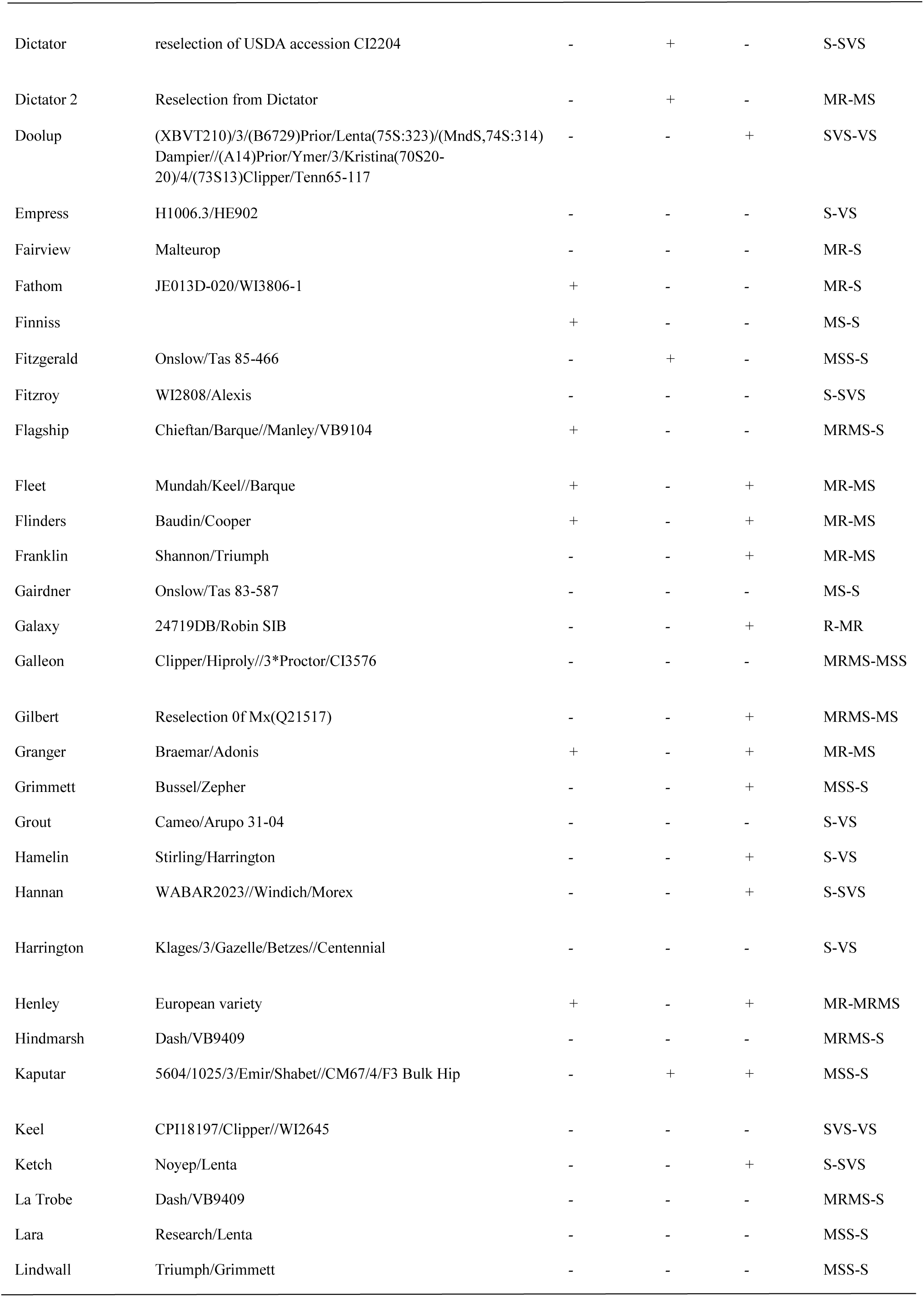
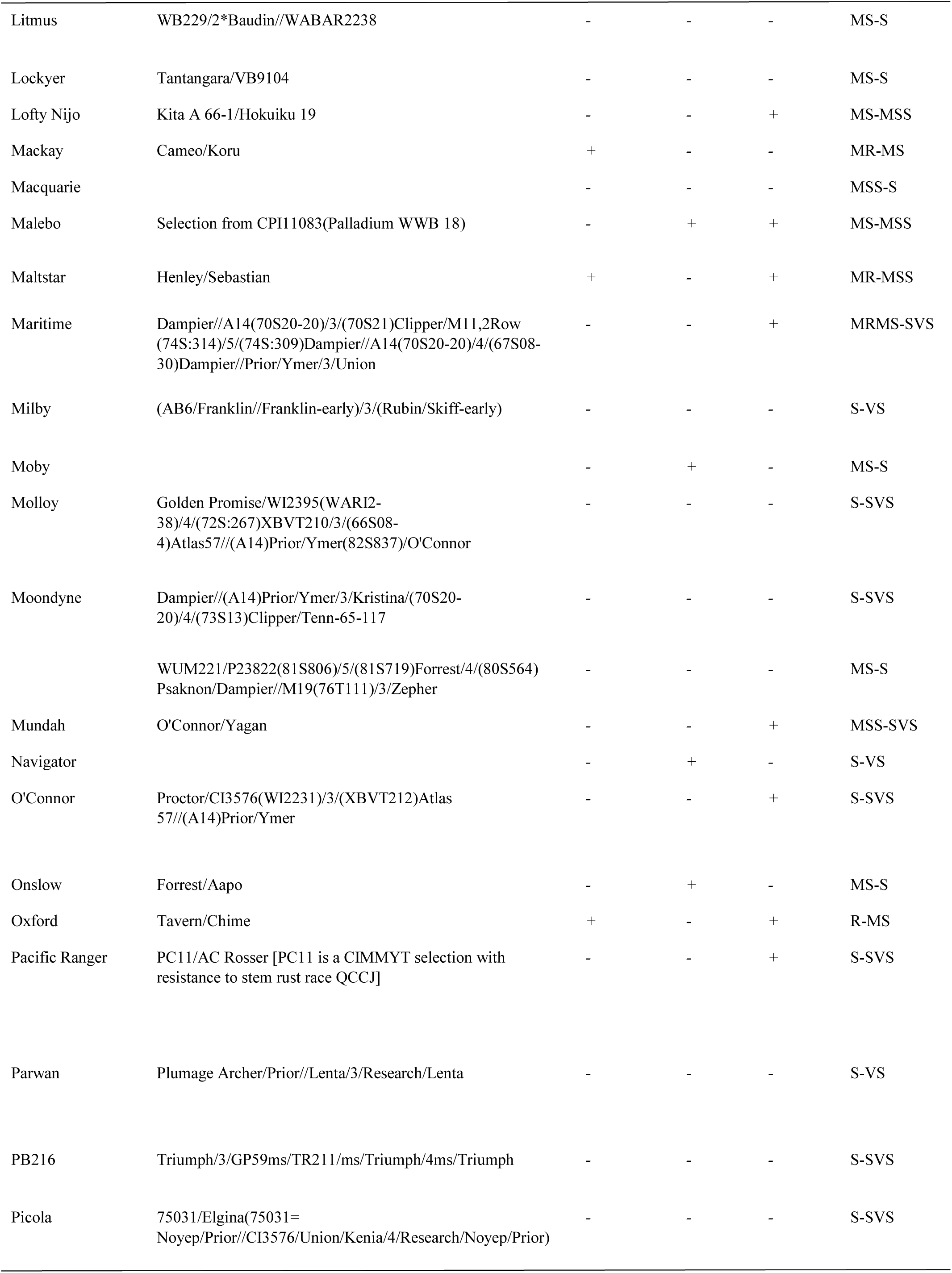
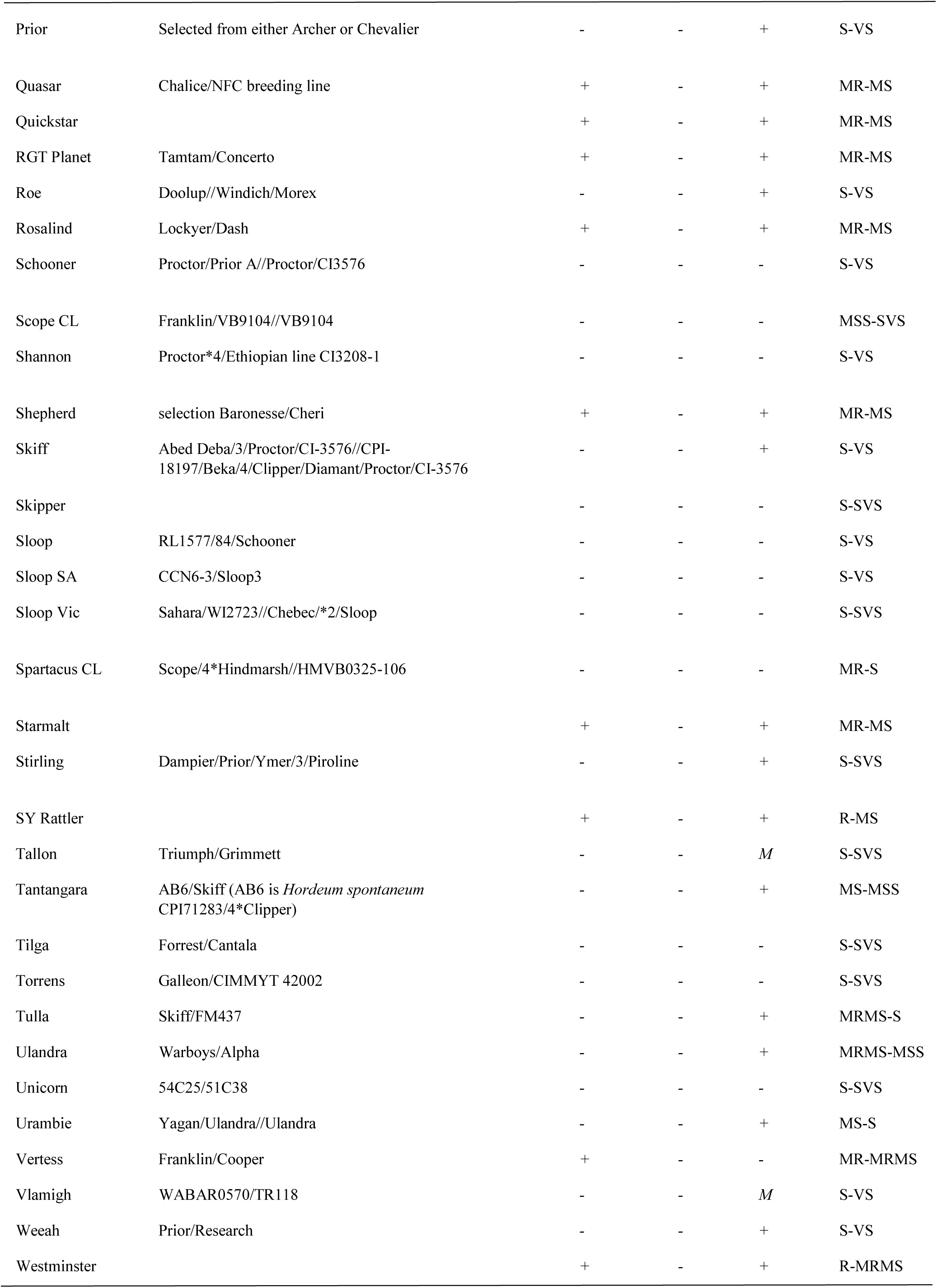
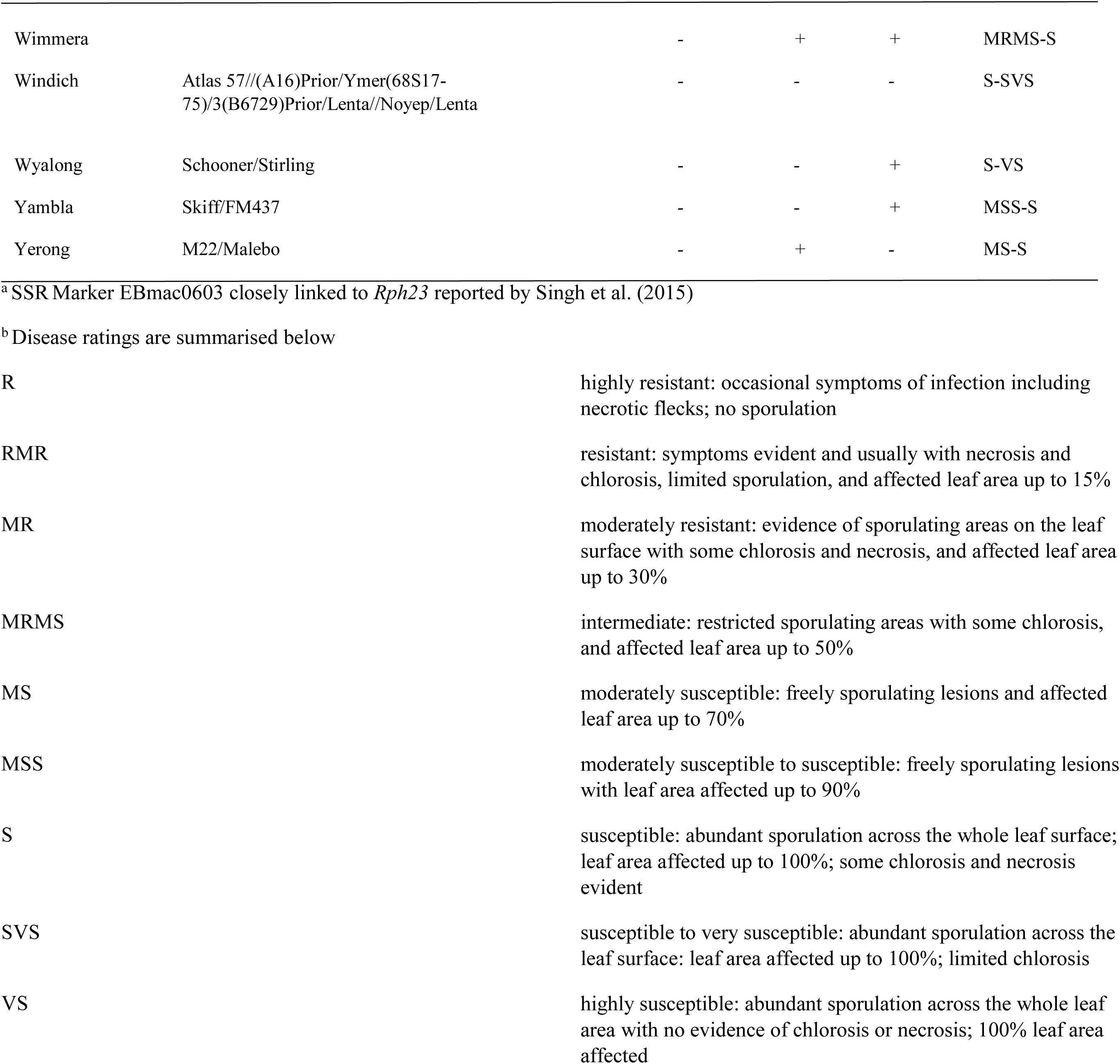
Summary information of postulated adult plant resistance status within 115 Australian barley cultivars based on the presence of co-dominant marker alleles associated with both resistance and susceptibility

| Cultivar | Pedigree | Adult Plant Resistance Gene |  |  | Consensus |
| --- | --- | --- | --- | --- | --- |
|  |  | *Rph20* | *Rph23<sup>a</sup>* | *Rph24* | Rating<sup>b</sup> |
| Arapiles | Noyep/Proctor//CI3576/Union/4/Kenia/3/Research/2/Noyep/Proctor/5/Domen | - | - | - | S-SVS |
| Bandulla | Prior/Lenta//Noyep/Lenta | - | - | - | S-SVS |
| Banks | WABAR2312/WABAR2332 | - | - | - | MS-S |
| Barque | Triumph/Galleon | + | - | - | MS-S |
| Bass | WABAR2023/Alexis | - | - | - | S-VS |
| Baudin | Stirling/Franklin | - | - | + | S-VS |
| Beecher | Atlas/Vaughan | - | + | + | MSS-S |
| Binalong | Blenheim//Skiff/O'Connor | - | - | + | S-VS |
| Bottler |  | + | - | + | R-MS |
| Brindabella | Weeah/CI7115//HCB27/3/Jadar II/4/Cantala | - | - | - | S-VS |
| Buloke | Franklin/VB9104//VB9104 | - | - | - | S-SVS |
| Bussell | Prior/Ymer | - | - | - | S-VS |
| Capstan | Waveney/WI2875//Chariot/Chebec | + | - | + | MR-MSS |
| Cask | SCRI-8313/Fleet//Regatta | - | - | + | MS-MSS |
| Charger |  | + | - | + | R-MRMS |
| Chebec | (USA) Orge Marin/2*Clipper(86)//Schooner (Aust.) (Orge Martin*Clipper#2)/86*Schooner | - | - | - | S-SVS |
| Chieftain | Brittania/Prisma | + | - | + | MR-MSS |
| Clipper | Proctor/Prior A | - | - | - | S-SVS |
| Commander | Keel/Sloop//Galaxy | - | - | - | MS-S |
| Compass | County/Commander//Commander | - | - | - | S-VS |
| Corvette | Bonus/CI3576 | + | - | - | MRMS-MSS |
| Cosmic |  | + | - | + | MR-MS |
| Cowabbie | (AB6/Franklin//Franklin-early)/3/(Rubin/Skiff-early) | - | - | + | S-SVS |
| Cutter | Proctor/Prior A | - | - | - | S-SVS |
| Dash | Chad/Joline//Cask | + | - | + | R-MRMS |
| Dhow | WI2808//Skiff/Haruna Nijo 9 | - | - | - | S-SVS |
| Dictator | reselection of USDA accession CI2204 | - | + | - | S-SVS |
| Dictator 2 | Reselection from Dictator | - | + | - | MR-MS |
| Doolup | (XBVT210)/3/(B6729)Prior/Lenta(75S:323)/(MndS,74S:314)<br>Dampier/(A14)Prior/Ymer/3/Kristina(70S20-20)/4/(73S13)Clipper/Tenn65-117 | - | - | + | SVS-VS |
| Empress | H1006.3/HE902 | - | - | - | S-VS |
| Fairview | Malteurop | - | - | - | MR-S |
| Fathom | JE013D-020/WI3806-1 | + | - | - | MR-S |
| Finniss |  | + | - | - | MS-S |
| Fitzgerald | Onslow/Tas 85-466 | - | + | - | MSS-S |
| Fitzroy | WI2808/Alexis | - | - | - | S-SVS |
| Flagship | Chieftan/Barque//Manley/VB9104 | + | - | - | MRMS-S |
| Fleet | Mundah/Keel//Barque | + | - | + | MR-MS |
| Flinders | Baudin/Cooper | + | - | + | MR-MS |
| Franklin | Shannon/Triumph | - | - | + | MR-MS |
| Gairdner | Onslow/Tas 83-587 | - | - | - | MS-S |
| Galaxy | 24719DB/Robin SIB | - | - | + | R-MR |
| Galleon | Clipper/Hiproly//3*Proctor/CI3576 | - | - | - | MRMS-MSS |
| Gilbert | Reselection Of Mx(Q21517) | - | - | + | MRMS-MS |
| Granger | Braemar/Adonis | + | - | + | MR-MS |
| Grimmett | Bussel/Zepher | - | - | + | MSS-S |
| Grout | Cameo/Arupo 31-04 | - | - | - | S-VS |
| Hamelin | Stirling/Harrington | - | - | + | S-VS |
| Hannan | WABAR2023//Windich/Morex | - | - | + | S-SVS |
| Harrington | Klages/3/Gazelle/Betzes//Centennial | - | - | - | S-VS |
| Henley | European variety | + | - | + | MR-MRMS |
| Hindmarsh | Dash/VB9409 | - | - | - | MRMS-S |
| Kaputar | 5604/1025/3/Emir/Shabet//CM67/4/F3 Bulk Hip | - | + | + | MSS-S |
| Keel | CPI18197/Clipper//WI2645 | - | - | - | SVS-VS |
| Ketch | Noyep/Lenta | - | - | + | S-SVS |
| La Trobe | Dash/VB9409 | - | - | - | MRMS-S |
| Lara | Research/Lenta | - | - | - | MSS-S |
| Lindwall | Triumph/Grimmett | - | - | - | MSS-S |
| Litmus | WB229/2*Baudin//WABAR2238 | - | - | - | MS-S |
| Lockyer | Tantangara/VB9104 | - | - | - | MS-S |
| Lofty Nijo | Kita A 66-1/Hokuiku 19 | - | - | + | MS-MSS |
| Mackay | Cameo/Koru | + | - | - | MR-MS |
| Macquarie |  | - | - | - | MSS-S |
| Malebo | Selection from CPII 1083(Palladium WWB 18) | - | + | + | MS-MSS |
| Maltstar | Henley/Sebastian | + | - | + | MR-MSS |
| Maritime | Dampier//A14(70S20-20)/3/(70S21)Clipper/M11,2Row<br>(74S:314)/5/(74S:309)Dampier//A14(70S20-20)/4/(67S08-30)Dampier//Prior/Ymer/3/Union | - | - | + | MRMS-SVS |
| Milby | (AB6/Franklin//Franklin-early)/3/(Rubin/Skiff-early) | - | - | - | S-VS |
| Moby |  | - | + | - | MS-S |
| Molloy | Golden Promise/WI2395(WARI2-38)/4/(72S:267)XBVT210/3/(66S08-4)Atlas57/(A14)Prior/Ymer(82S837)/O'Connor | - | - | - | S-SVS |
| Moondyne | Dampier//A14)Prior/Ymer/3/Kristina/(70S20-20)/4/(73S13)Clipper/Tenn-65-117 | - | - | - | S-SVS |
|  | WUM221/P23822(81S806)/5/(81S719)Forrest/4/(80S564)Psaknon/Dampier//M19(76T111)/3/Zepher | - | - | - | MS-S |
| Mundah | O'Connor/Yagan | - | - | + | MSS-SVS |
| Navigator |  | - | + | - | S-VS |
| O'Connor | Proctor/CI3576(WI2231)/3/(XBVT212)Atlas57/(A14)Prior/Ymer | - | - | + | S-SVS |
| Onslow | Forrest/Aapo | - | + | - | MS-S |
| Oxford | Tavern/Chime | + | - | + | R-MS |
| Pacific Ranger | PC11/AC Rosser [PC11 is a CIMMYT selection with resistance to stem rust race QCCJ] | - | - | + | S-SVS |
| Parwan | Plumage Archer/Prior//Lenta/3/Research/Lenta | - | - | - | S-VS |
| PB216 | Triumph/3/GP59ms/TR211/ms/Triumph/4ms/Triumph | - | - | - | S-SVS |
| Picola | 75031/Elgina(75031=Noyep/Prior//CI3576/Union/Kenia/4/Research/Noyep/Prior) | - | - | - | S-SVS |
| Prior | Selected from either Archer or Chevalier | - | - | + | S-VS |
| Quasar | Chalice/NFC breeding line | + | - | + | MR-MS |
| Quickstar |  | + | - | + | MR-MS |
| RGT Planet | Tamtam/Concerto | + | - | + | MR-MS |
| Roe | Doolup//Windich/Morex | - | - | + | S-VS |
| Rosalind | Lockyer/Dash | + | - | + | MR-MS |
| Schooner | Proctor/Prior A//Proctor/CI3576 | - | - | - | S-VS |
| Scope CL | Franklin/VB9104//VB9104 | - | - | - | MSS-SVS |
| Shannon | Proctor*4/Ethiopian line CI3208-1 | - | - | - | S-VS |
| Shepherd | selection Baronesse/Cheri | + | - | + | MR-MS |
| Skiff | Abed Deba/3/Proctor/CI-3576//CPI-18197/Beka/4/Clipper/Diamant/Proctor/CI-3576 | - | - | + | S-VS |
| Skipper |  | - | - | - | S-SVS |
| Sloop | RL1577/84/Schooner | - | - | - | S-VS |
| Sloop SA | CCN6-3/Sloop3 | - | - | - | S-VS |
| Sloop Vic | Sahara/WI2723//Chebec/*2/Sloop | - | - | - | S-SVS |
| Spartacus CL | Scope/4*Hindmarsh//HMBV0325-106 | - | - | - | MR-S |
| Starmalt |  | + | - | + | MR-MS |
| Stirling | Dampier/Prior/Ymer/3/Piroline | - | - | + | S-SVS |
| SY Rattler |  | + | - | + | R-MS |
| Tallon | Triumph/Grimmett | - | - | *M* | S-SVS |
| Tantangara | AB6/Skiff (AB6 is *Hordeum spontaneum* CPI71283/4*Clipper) | - | - | + | MS-MSS |
| Tilga | Forrest/Cantala | - | - | - | S-SVS |
| Torrens | Galleon/CIMMYT 42002 | - | - | - | S-SVS |
| Tulla | Skiff/FM437 | - | - | + | MRMS-S |
| Ulandra | Warboys/Alpha | - | - | + | MRMS-MSS |
| Unicorn | 54C25/51C38 | - | - | - | S-SVS |
| Urambie | Yagan/Ulandra//Ulandra | - | - | + | MS-S |
| Vertess | Franklin/Cooper | + | - | - | MR-MRMS |
| Vlamigh | WABAR0570/TR118 | - | - | *M* | S-VS |
| Weeah | Prior/Research | - | - | + | S-VS |
| Westminster |  | + | - | + | R-MRMS |
| Wimmera |  | - | + | + | MRMS-S |
| Windich | Atlas 57/(A16)Prior/Ymer(68S17-75)/3(B6729)Prior/Lenta//Noyep/Lenta | - | - | - | S-SVS |
| Wyalong | Schooner/Stirling | - | - | + | S-VS |
| Yambla | Skiff/FM437 | - | - | + | MSS-S |
| Yerong | M22/Malebo | - | + | - | MS-S |

<sup>a</sup> SSR Marker EBmac0603 closely linked to *Rph23* reported by Singh et al. (2015)
<sup>b</sup> Disease ratings are summarised below

|  |  |
| --- | --- |
| R | highly resistant: occasional symptoms of infection including necrotic flecks; no sporulation |
| RMR | resistant: symptoms evident and usually with necrosis and chlorosis, limited sporulation, and affected leaf area up to 15% |
| MR | moderately resistant: evidence of sporulating areas on the leaf surface with some chlorosis and necrosis, and affected leaf area up to 30% |
| MRMS | intermediate: restricted sporulating areas with some chlorosis, and affected leaf area up to 50% |
| MS | moderately susceptible: freely sporulating lesions and affected leaf area up to 70% |
| MSS | moderately susceptible to susceptible: freely sporulating lesions with leaf area affected up to 90% |
| S | susceptible: abundant sporulation across the whole leaf surface; leaf area affected up to 100%; some chlorosis and necrosis evident |
| SVS | susceptible to very susceptible: abundant sporulation across the leaf surface: leaf area affected up to 100%; limited chlorosis |
| VS | highly susceptible: abundant sporulation across the whole leaf area with no evidence of chlorosis or necrosis; 100% leaf area affected |

**Figure 2.**
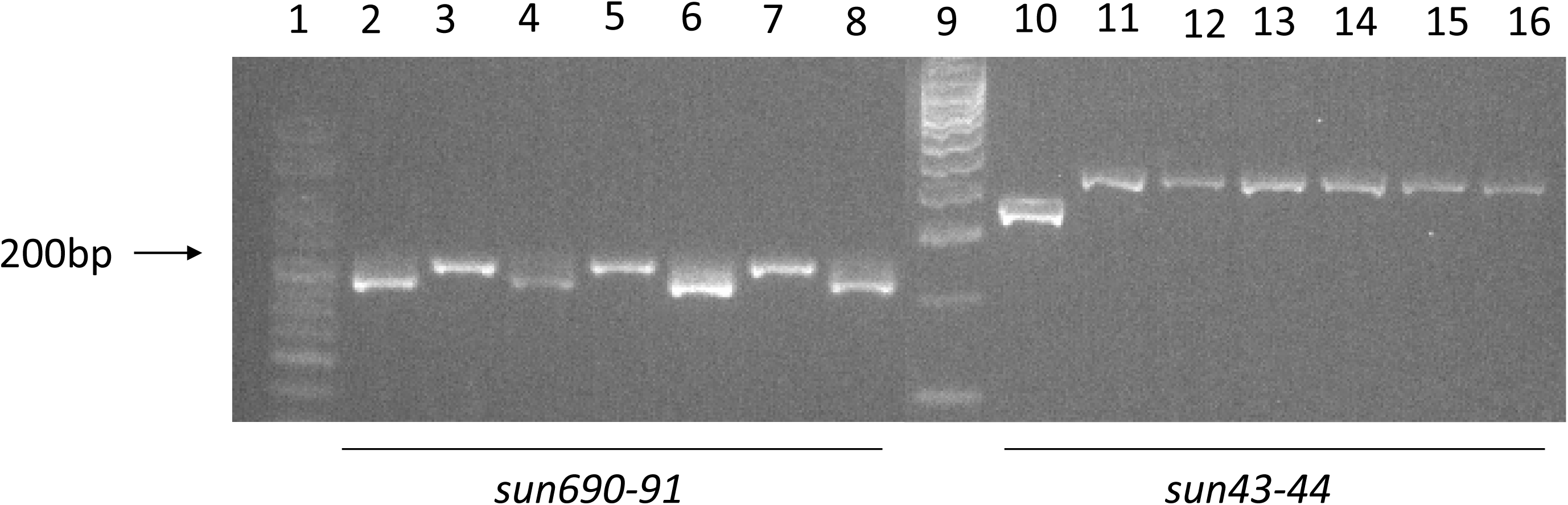
Results of marker genotypic analysis on eight barley cultivars with co-dominant markers linked to adult plant resistance (APR) genes *Rph20* (*sun690-91*) and *Rph24* (*sun43-44*). Three controls were used including: Flagship (*Rph20/rph24*) [2 and 10], ND24260 (*Rph24/rph20*) [3 and 11], and Vada (*Rphq3/Rphq4*) [4 and 12] were run alongside four Australian cultivars carrying varying levels of APR including Grimmet [5 and 13], Westminister [6 and 14], Maritime [7 and 15] and Oxford [8 and 16]. PCR products were separated on a 2.5% agarose gel and the size was determined using Bioline hyperladders 25bp (1) and 100bp (9).

To date we have genotyped all 320 accessions from the association mapping panel reported in Singh et al. (2018) with the three markers (data not shown). Marker data for both *Rph20* markers including the dominant bPb-0837 (Hickey et al. 2011) and the co-dominant indel marker developed in the present study correlated completely, suggesting both are strongly predictive for the *Rph20* resistance. Our marker data corroborates previous findings that *Rph20* and *Rph24* are found in combination at a high frequency (Zeims et al. 2017; Singh et al. 2018). Previous studies suggest that *Rph20* is not present or detected at very low frequencies in germplasm derived from China (Derevnina et al. 2013) and Africa (Elmansour et al. 2017), however the *Rph24* marker allele was identified in more than 50% of these accessions. This suggests that *Rph24* may be more widespread globally relative to the *Rph20* conferred resistance. Within European lines, German cultivar Volla and Lenka carried all three APR genes and the all stage resistance gene *Rph3* and had a consensus adult plant rating of R-MR. For bread wheat, near-immunity in the field has previously been attributed to the presence of multiple APR genes in combination (Lillemo et al. 2011). The pathotype of *P. hordei* used in the Cobbitty field nursery leaf rust resistance plots carried virulence for the *Rph3* resistance (Park RF. unpublished) however the combination of all three APR genes provided immunity to leaf rust despite the high disease severity in the majority of year assessed. Other European lines including RAH1995, Klimek and Baronesse carried the *Rph20*/*Rph24* combination. In contrast, Australian cultivars Beecher, Malebo and Kaputar were found to carry both *Rph23* and *Rph24* despite developing significant levels of leaf rust at adult plant growth stages in the field especially under high disease pressure. Hence, the combination of *Rph23* and *Rph24* does not appear to provide adequate protection.

In conclusion, *Rph20* and *Rph24* are both widely distributed genes conferring APR to *P. hordei* that when deployed in combination offer high levels of protection. *Rph20* offers barley growers the most protection and has remained effective to all known variants of *P. hordei* despite its prevalence in global germplasm and hence widespread deployment. Here, we report on the development and validation of co-dominant PCR-based markers to improve the efficiency of MAS for APR to leaf rust resistance. The development of co-dominant markers for all three characterised APR genes described to date for BLR has permitted the diversification of disease resistance through prioritising accessions that carry APR but lack all three marker alleles. In contrast, genotypic analysis also identified marker-positive accessions previously postulated carry uncharacterised APR that was likely explained by the presence of *Rph24*. Future work will focus on characterising new sources of APR in diverse barley accessions that lack all three known APR marker alleles.

## Acknowledgements

The authors would like to thank the Grains Research and Development Corporation for funding the research under project US00074 and Ms Bethany Clark for technical support.

